# Assessing the remarkable morphological diversity and transcriptomic basis of leaf shape in *Ipomoea batatas* (sweetpotato)

**DOI:** 10.1101/520650

**Authors:** Sonal Gupta, David M. Rosenthal, John R. Stinchcombe, Regina S. Baucom

## Abstract

- Leaf shape, a spectacularly diverse plant trait, varies across taxonomic levels, geography, and in response to environmental differences. However, comprehensive intraspecific analyses of leaf shape variation across variable environments is surprisingly absent. Here, we perform a multi-level analysis of leaf shape using diverse accessions of sweetpotato (*Ipomoea batatas*), and uncover the role of genetics, environment, and GxE on this important trait.
- We examine leaf shape using a variety of morphometric analyses, and complement this with a transcriptomic survey to identify gene expression changes associated with shape variation. Additionally, we examine the role of genetics and environment on leaf shape by performing field studies in two geographically separate common gardens.
- We show that extensive leaf shape variation exists within *I. batatas*, and identify promising candidate genes underlying this variation. Interestingly, when considering traditional measures, we find that genetic factors are largely responsible for most of leaf shape variation, but that the environment is highly influential when using more quantitative measures *via* leaf outlines.
- This extensive and multi-level examination of leaf shape shows an important role of genetics underlying a potentially important agronomic trait, and highlights that the environment can be a strong influence when using more quantitative measures of leaf shape.

## INTRODUCTION

Leaf shape varies spectacularly among plant species at multiple taxonomic levels (Klein *et al*., 2017; Shi *et al*., 2019), across geography (Wyatt & Antonovics, 1981; Gurevitch, 1988), and in response to environmental differences (Andersson, 1991; Jones, 1995; McDonald *et al*., 2003). Leaves can vary with respect to their degree of dissection, length-to-width ratio, venation patterning, prominence of tips and petiolar sinus, or any combinations of the above, meaning that leaf shape variation across species is multifaceted and complex. Leaf shape diversity is also present within species (Hilu, 1983). For example, accessions of grapevine and cotton vary with respect to leaf complexity whereas lineages within tomato and apple show ample variation in the length-to-width ratio of leaves (Chitwood *et al*., 2013; Andres *et al*., 2016; Klein *et al*., 2017; Migicovsky *et al*., 2017). Although a large number of species exhibit variation in leaf shape, examinations within species are often limited to only a few accessions, with a few notable exceptions (Conesa *et al*., 2012; Chitwood *et al*., 2014a, b). Moreover, these studies often focus on circularity and length-to-width ratio, which are the most common leaf shape descriptors. Thus, for most species, truly quantitative analyses of the diversity of leaf shape variation within species remains largely unexamined.

Leaf shape variation is regulated by genetics, the environment, and the interaction of genes and environment (GxE). Although the genetic and trancriptomic basis underlying leaf shape diversity has been uncovered in only a small number of species *(i.e.*, tomato, Arabidopsis, cotton, and a few others; Kim *et al.*, 2002; Kimura *et al*., 2008; Vlad *et al*., 2014; Ichihashi *et al*., 2014; Andres *et al*., 2016; Chitwood & Sinha, 2016), there are many examples showing the influence of different environments on leaf shape (McDonald *et al*., 2003; Zwieniecki *et al*., 2004; Hopkins *et al*., 2008; Royer *et al*., 2009; Nicotra *et al*., 2011; Royer, 2012; Campitelli & Stinchcombe, 2013; Glennon & Cron, 2015). For example, submerged leaves of aquatic plants are often highly dissected as compared to their aerial counterparts (Arber, 2010) and leaves growing in colder environments tend to be more complex than similar ones growing in warmer environments (Huff *et al*., 2003; Royer *et al*., 2005). Moreover, the environment can interact with genes to further modulate leaf shape. For instance, Nakayama and colleagues (2014) found that changes in temperature leads to abrupt changes in KNOX1 (*KNOTTED1-LIKE HOMOEOBOX1*) activity, a key regulator of circularity in multiple species, thus altering leaf complexity. Although we are beginning to understand how genetics, environment, and GxE separately influence aspects of leaf shape, few studies have partitioned the effect of genetics versus the environment on leaf shape variation, and most examinations are limited to only one environment, such that the role of GxE on leaf shape is often not considered within species.

Leaf shape is most commonly quantified using the ‘traditional’ leaf shape traits – circularity (a measure of leaf dissection, or ‘lobedness’), aspect ratio (the length-to-width ratio of a leaf) and solidity (the relation of the area and convex hull). These traditional morphometric parameters have previously been used to quantity leaf shape in diverse species, such as grapes (Chitwood *et al*., 2014b), tomato (Chitwood *et al*., 2015) and sweetpotato (Rosero *et al*., 2019), among others. Although these traits are linked to important yield traits in crops (Chitwood *et al*., 2013; Vuolo *et al*., 2016; Chitwood & Otoni, 2017; Klein *et al*., 2017; Rowland *et al*., 2019), and are important for understanding the broader aspects of plant adaptation to environment, they capture only a few components of leaf shape variation. A more comprehensive quantification of leaf shape can be captured with Elliptical Fourier Descriptor (EFD) analyses, which converts leaf outlines to harmonic coefficients allowing for Fourier analyses (Chitwood & Sinha, 2016). This approach captures extensive leaf shape variation due to both symmetry and asymmetry of the leaf; some examples include shape differences associated with the depth of the petiolar sinus, the prominence of the leaf tip, and the positioning of the lobes. This approach has been applied to a handful of species like tomatoes, passiflora, and grape (Chitwood *et al*., 2013; Chitwood & Otoni, 2017; Klein *et al*., 2017), where it was shown that leaf shape based on EFD analysis is highly heritable. Thus, traditional measures along with consideration of leaf outlines holds greater power to comprehensively measure and characterize leaf shape, which may yield important insights about the genetic basis of leaf shape variation. Interestingly, while leaf shape based on EFD analysis is heritable, no studies have yet examined the genetic or transcriptomic basis of leaf shape based on leaf outlines.

*Ipomoea batatas*, the sweetpotato, is an important staple root crop worldwide (Khoury *et al*., 2015), as it produces the highest amount of edible energy per hectare (Khoury *et al*., 2015) and also provides an important source of nutrients in the form of vitamin A, calcium, and iron (Kays & Kays, 1998). Sweetpotato displays striking morphological variation in leaf shape across its ~6000 documented varieties (Huaman, 1987), but very few studies have examined the extensive leaf shape diversity in this species (Huaman, 1987; Hue *et al*., 2012; Rosero *et al*., 2019). Studies that have examined leaf shape phenotypes in sweetpotato are limited to a few cultivars and/or present traditional measures of leaf shape traits. Additionally, the genetic or transcriptomic basis of leaf shape variation in this species has yet to be considered. The vast unexamined diversity of leaf shape in this species, along with its role as a staple food crop worldwide makes *I. batatas* an ideal study system to investigate leaf shape diversity at the species level and how this diversity is influenced by the interplay between genetics and environment.

Here, we examine the extensive leaf shape variation within accessions of *I. batatas*, and uncover the role of genetics, environment and GxE in influencing leaf shape traits. We specifically ask: (1) How diverse is leaf shape at a species-wide level? (2) what are the candidate genes associated with leaf shape (extending beyond the traditional shape descriptors)? and (3) to what degree does the environment and GxE influence leaf shape traits? We show that extensive natural variation exists in leaf shape within this species and that most of this variation is largely controlled by genetic factors, with a low proportion of variance in leaf shape attributable to environmental differences. We also identified promising candidate genes that underlie broad differences in multiple leaf shape traits. The results of our work fill critical gaps in current knowledge of leaf shape evolution by expanding analysis beyond that of the traditional measures of leaf shape and by using many distinct lineages of the species. We unite this with the transcriptomic basis of these traits along with a multiple-environment assessment of leaf shape variation in the field. Thus, this work allows us to comprehensively assess leaf shape in this agronomically important species and partition the role of genetics, environment, and GxE on leaf shape within this species.

## METHODS

### Leaf shape variation within *I. batatas*

We ordered vegetative slips for 68 publicly available accessions of sweetpotato from USDA and online resources. The location of origin of 68 accessions is represented in Fig. 1 (Table S1). The accessions represent the majority of the genetic variation in the species; we identified three of the four population structure clusters among our chosen accessions as per a recent study (Wadl *et al*., 2018). We grew slips at the UM Matthaei Botanical Garden under standardized growth conditions (16 hrs light/8 hrs night cycle) for approximately six months, at which time we sampled 4-6 mature leaves (third-sixth mature leaves from the beginning of the vine to control for age and exposure to light) of 57 randomly chosen accessions and scanned them for leaf shape analyses.

**Figure 1.**
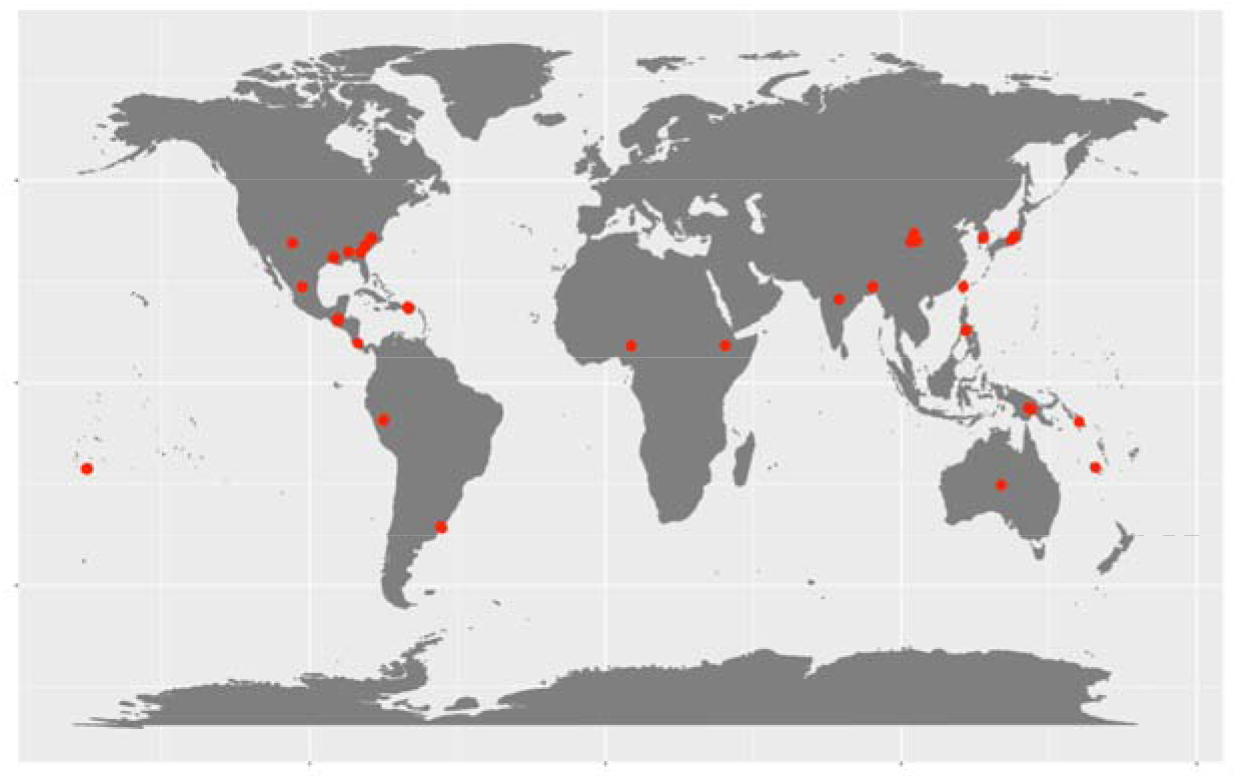
Geographic diversity of the 74 chosen sweetpotato, *Ipomoea batatas*, accessions. Red dots represent the origin of the chosen samples.

We used the scanned images to extract leaf shape trait values using custom macros in ImageJ (Abràmoff *et al*., 2004). Briefly, we converted leaves into binary images and then used outlines from these binary images to measure circularity, aspect ratio and solidity, each capturing a distinct aspect of leaf shape (Li *et al*., 2018). Circularity, measured as 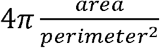, is influenced by serrations and lobing. Aspect ratio, in comparison, is measured as the ratio of the major axis to the minor axis of the best fitted ellipse, and is influenced by leaf length and width. Lastly, solidity measured as 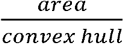, is sensitive to leaves with deep lobes, or with a distinct petiole, and can be used to distinguish leaves lacking such structures. Solidity, unlike circularity, is not very sensitive to serrations and minor lobings, since the convex hull remains largely unaffected.

For a more global analysis of leaf shape via Elliptical Fourier Descriptor (EFDs), we used the program SHAPE (Iwata & Ukai, 2002) as described in (Chitwood *et al*., 2014b). EFDs capture variation in shape represented by the outline which is difficult to categorize via traditional shape descriptors. From the EFD coefficients obtained, we used coefficients a and d only, thus analyzing symmetric variation in leaf shape. Principal component analysis (PCA) was performed on the EFD coefficients to identify shape features contributing to leaf morphological variation (referred to as EFD symPCs below). We calculated the correlation matrices using the rcorr() function of the Hmisc package version 4.0-3 (Harrell *et al*., 2017) with multiple test adjustments using the p.adjust() function in R.

### RNA-Seq library construction and sequencing

We sequenced and analyzed transcriptomes of 19 individuals of *I. batatas* to examine gene expression differences associated with leaf shape variation associated with circularity, aspect ratio, and EFD symPCs to obtain an initial set of candidate genes underlying these traits. We selected greenhouse-grown accessions with differing leaf shape trait values (Fig. S1). Since high aspect ratio represents both longitudinally longer or latitudinally broader leaf shape phenotypes, we chose to only examine individuals that had high aspect ratio due to latitudinal elongation. We chose multiple accessions to assess each leaf shape trait; eleven for circularity (six entire, five lobed), eight for aspect ratio (four high and low AR, each), 6 individuals for EFD symPC1 (three high and three low) and four accessions each for EFD symPC2 and EFD symPC3 (two high and two low) (Fig. S1); EFD symPC4 was not considered for differential expression analysis.

We used three to five leaves that were in P4-P6 stage of growth (fourth to sixth youngest primordium), from multiple branches of each individual accession for RNA extractions, and combined replicate leaves per individual to increase the depth of the transcriptome. We sampled all individuals on the same day within 1 hour to reduce variation due to developmental stage and/or time of collection. We froze samples in liquid nitrogen prior to preserving them at −80° for further processing. We performed RNA extraction using Qiagen RNeasy Plant mini kit with the optional DNase digestion step, and constructed libraries using the TruSeq Stranded mRNA Sample Preparation protocol (LS protocol). After barcoding, we bulked all libraries and performed one lane of Illumina HiSeq2500 sequencing.

### RNA-Seq data processing and transcriptome analysis

An overview of our RNA-Seq data processing and transcriptome analysis is given in Fig. 2, with detailed information presented in Method S1.

**Figure 2.**
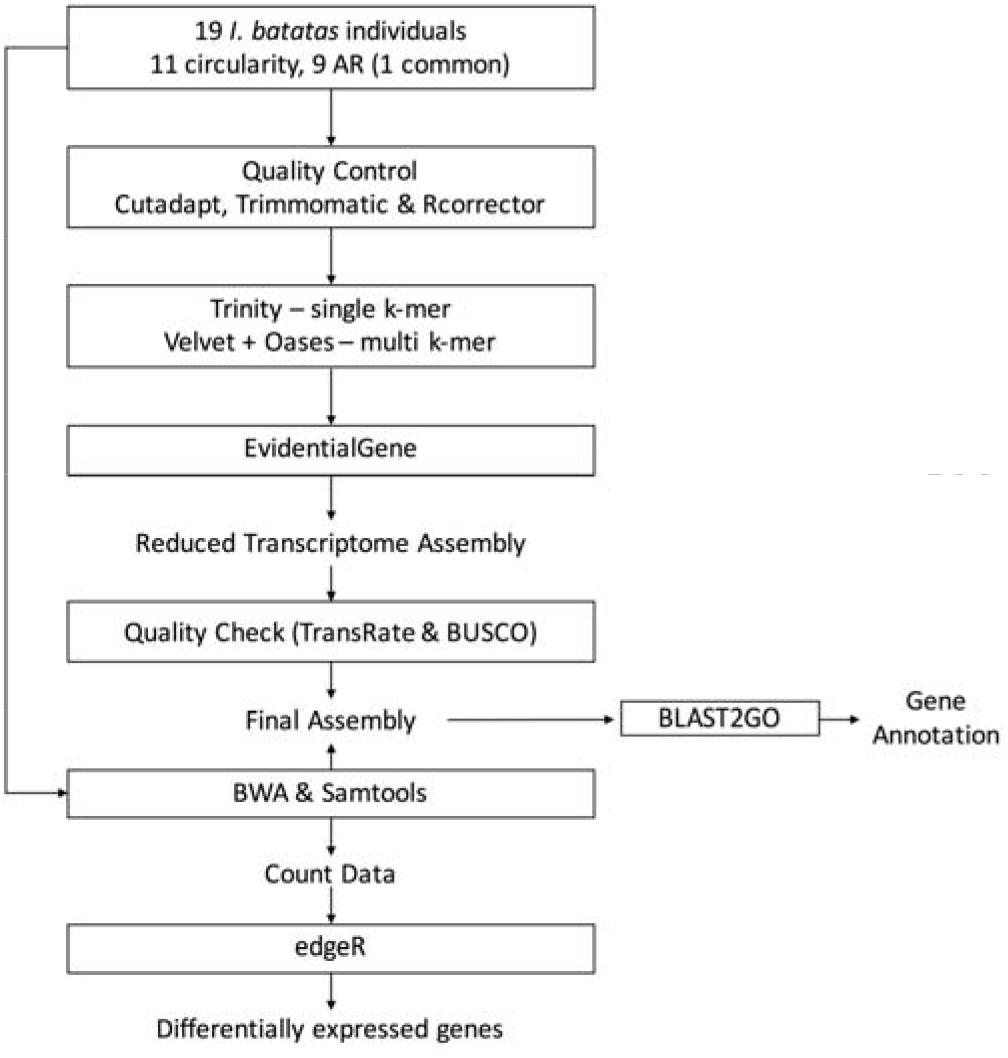
Methodology for RNA-Seq data processing for differential gene expression

#### Differential gene expression

We mapped reads from all 19 individuals to the *de novo* assembled transcriptome using BWA-MEM v0.7.15 (Li, 2013) and estimated read counts for uniquely mapped reads using samtools v1.9 (Li *et al*., 2009). We then used read counts to filter out lowly expressed transcripts using the Bioconductor package edgeR version 3.18.1 (Robinson *et al*., 2010) such that transcripts were retained only if they had greater than 0.5 counts-per-million in at least two samples. We then normalized libraries in edgeR (using the trimmed mean of *M*-values method) followed by differential gene expression analysis using classic pairwise comparison of edgeR version 3.18.1. We extracted the significance of differentially expressed transcripts (DETs) with FDR <= 0.05.

### Field experiment

We performed a field experiment to determine the extent to which genetics, the environment, and GxE interactions influence leaf shape traits. We generated replicate individuals by planting 5 cm cuttings of the stem of each accession in 4-inch pots, randomly positioned on a mist bench at the Matthaei Botanical Gardens. During the first week of June, we planted three to seven replicates of each of the 68 accessions in two common gardens--one located at the Matthaei Botanical Gardens in Ann Arbor, MI (42.18° N, 83.39° W), and the other at the Ohio University Student Farm, West State Street Research Site in Athens, OH (40.46° N, 81.55° W). Replicates were planted in either three (MI) or seven (OH) blocks in a completely randomized block design with 14-inch spacing between individuals. Blocks were kept relatively weed free but were otherwise allowed to grow undisturbed. We randomly sampled 2-5 mature leaves from each individual in the first week of October, prior to the first frost, and scanned them for leaf shape analyses as explained before.

#### Data analysis

We first examined the potential for variation in leaf shape due to environmental differences (i.e. variation due to being grown in MI or OH) by performing an ANOVA. To normalize leaf shape traits, we used the function TransformTukey from rcompanion version 2.0.0 (Mangiafico, 2018). TransformTukey is a power transformation based on Tukey’s ladder of Powers, which loops through multiple powers and selects the one that normalizes the data most. These normalized leaf shape traits were then used as dependent variables and accession, garden, block effects and an interaction term of accession and garden as independent variables in the following fixed-effects model:

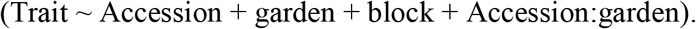

The term accession represents the genetic component, garden represents variation due to environment (plasticity), Accession:garden represents the GxE component and the block effect captures microenvironmental variation (and was nested within each garden). To quantify the relative effects of each of these variables on leaf shape, we calculated eta squared (η2) as a measure of the magnitude of effect size using the Bioconductor package lsr version 0.5 (Navarro, 2013). Eta squared for an effect is measured as SS_effect_/SS_total_, where SS_effect_ is the sum of squares of the effect of interest and SS_total_ is the total sum of squares of all the effects, including interactions. In other words, it is a measure of the proportion of variance in the dependent variable associated with independent variable and is one of the most commonly reported estimates of effect size for ANOVA (Levine & Hullett, 2002; Ialongo, 2016). Further, we calculated broad sense heritabilities of leaf shape traits to determine the extent to which traits are genetically controlled within each environment. Broad sense heritability was calculated using linear mixed modeling with the Bioconductor package sommer version 3.4 (Covarrubias-Pazaran, 2016) based on the phenotypic data collected from the two fields. The model used was

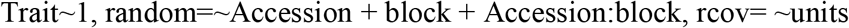

Variance components from the model were used to calculate the broad-sense heritability (H^2^) using the formula:

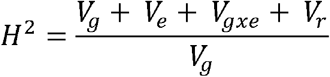

where V_g_ is the genotype variance, V_e_ is the environmental variance due to the blocks, V_gxe_ is the variance associated with V_gxe_ (accession:block), and V_r_ is the residual variance.

## RESULTS

### Leaf shape variation among accessions

We found wide variation in leaf traits across 57 *I. batatas* accessions (Table 1). Among the three traditional traits examined, circularity is most variable with a phenotypic coefficient of variation (PCV; (standard deviation(x)/mean(x))*100; where x is the trait of interest) of 22.61% while aspect ratio is least variable with a narrow distribution and PCV of 4.76%. Figure 3 shows the phenotypic diversity with respect to two leaf traits, circularity and aspect ratio (AR). Of our 57 accessions, 10 exhibit low circularity (defined as circularity < 0.50). PI 599387, for example, exhibited leaves that are very deeply lobed and thus has a low circularity (0.09) value. In contrast, PI 566647 has no serrations or lobing (entire margins) and thus exhibits high circularity (0.71; Fig. 3). Additionally, we found 22 of 57 accessions to exhibit high aspect ratio (AR > 1.11). For example, PI 531134 (AR = 1.03) has almost equal values of major and minor axis and thus a low aspect ratio value. In contrast, the leaves of PI 208886 (AR = 1.268) are much wider, i.e., a larger major to minor axis, and thus has high aspect ratio value. Most often this increase in AR in sweetpotato manifests itself with increase leaf width (eg. PI 566646, PI 208886) relative to length (eg. PI 634379). Further, although solidity values range from 0.44-0.95, only 5 accessions had solidity values less than 0.7 (PCV = 11.85%). The lack of low solidity values indicates that only a few accessions have deeply lobed leaves (eg. PI 599387, solidity = 0.44), in contrast to accessions with slightly lobed leaved (eg. PI 566630, solidity = 0.76).

**Figure 3.**
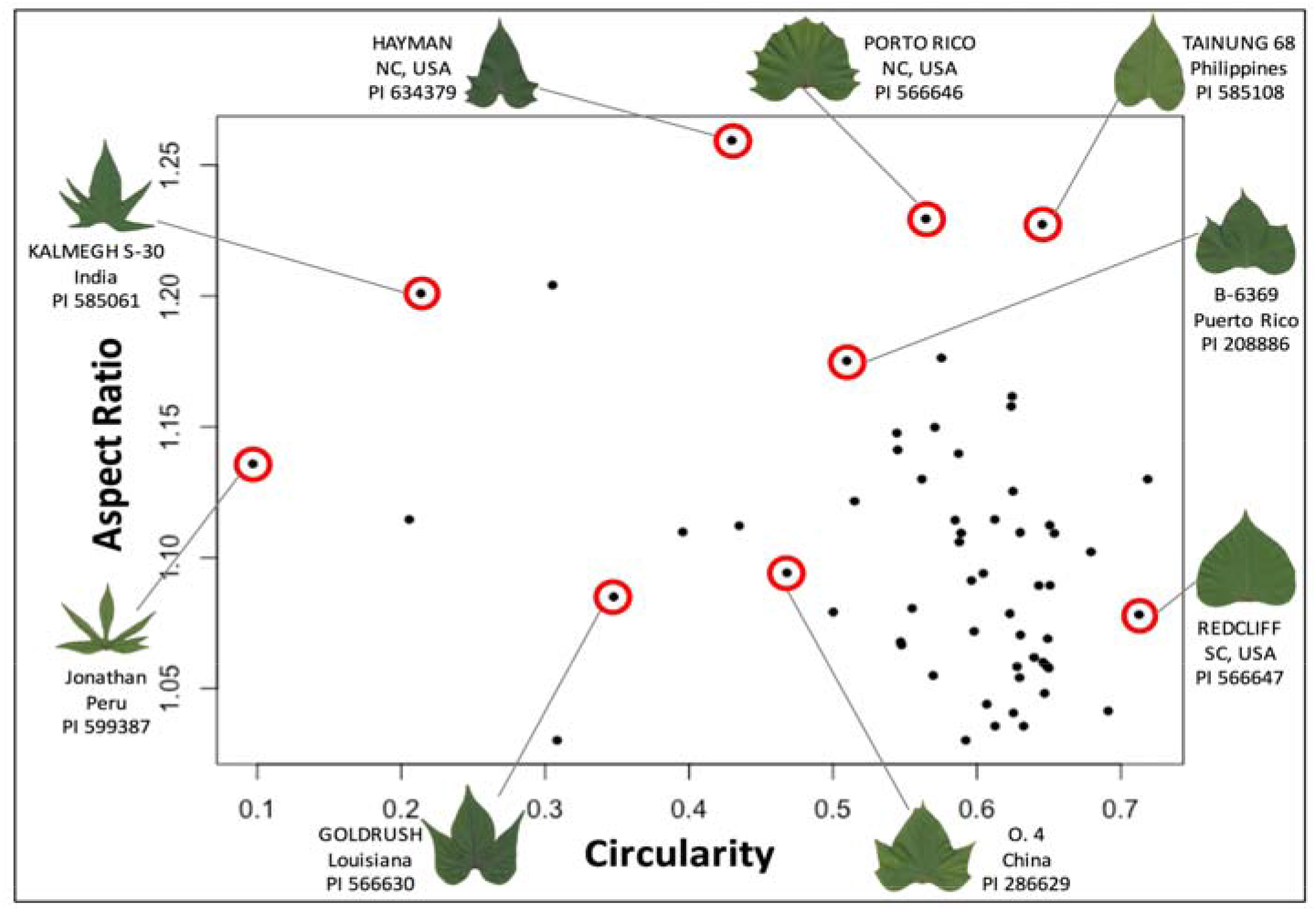
Leaf shape variation in a sample of accessions of sweetpotato, *Ipomoea batatas*, highlighting exceptionally high morphological variation.

**Table 1.** Leaf shape trait values across the 57 chosen sweetpotato accessions. SD represents standard deviation while PCV represents phenotypic coefficient of variation.

| Trait | Range | Mean | SD | PCV (%) |
| --- | --- | --- | --- | --- |
| Circularity | 0.09-0.71 | 0.50 | 0.12 | 22.61 |
| Aspect Ratio | 1.03-1.26 | 1.10 | 0.05 | 4.76 |
| Solidity | 0.44-0.95 | 0.84 | 0.10 | 11.85 |

We performed an EFD analysis on leaf outlines to get a more global estimation of leaf shape variation (Fig. 4). In total, we processed 292 leaves from 57 accessions to identify leaf shape traits that explain symmetrical shape variation in sweetpotato. Low symPC1 values describe leaves with deep lobing, prominent tip and shallow petiolar sinus (PI 573318) whereas high symPC1 values explain non-lobed leaves with flattened leaf tips and enclosed petiolar sinus (PI 566646). symPC2 explains variation in leaf shape due to differences in breadth and lobing of the leaf (low symPC2 values describe broad leaves with two lobes whereas high symPC2 values depicts narrow leaves with no lobes). symPC3 primarily captures leaf shape variation due to the depth of petiolar sinus (low symPC3 values describe leaves with highly enclosed petiolar sinus as compared to high symPC3 eigenleaves which have flattened sinus). Lastly, symPC4 represents variation in leaf shape attributed to the angle of lobe tips -- low symPC4 eigenleaves have lobes with a high obtuse angle (almost 160°) whereas high symPC4 eigenleaves have lobes with a lower obtuse angle (almost 125°). The four symPC components together explain 87.79% of total variance relating to symmetrical leaf shape variance in sweetpotato.

**Figure 4:**
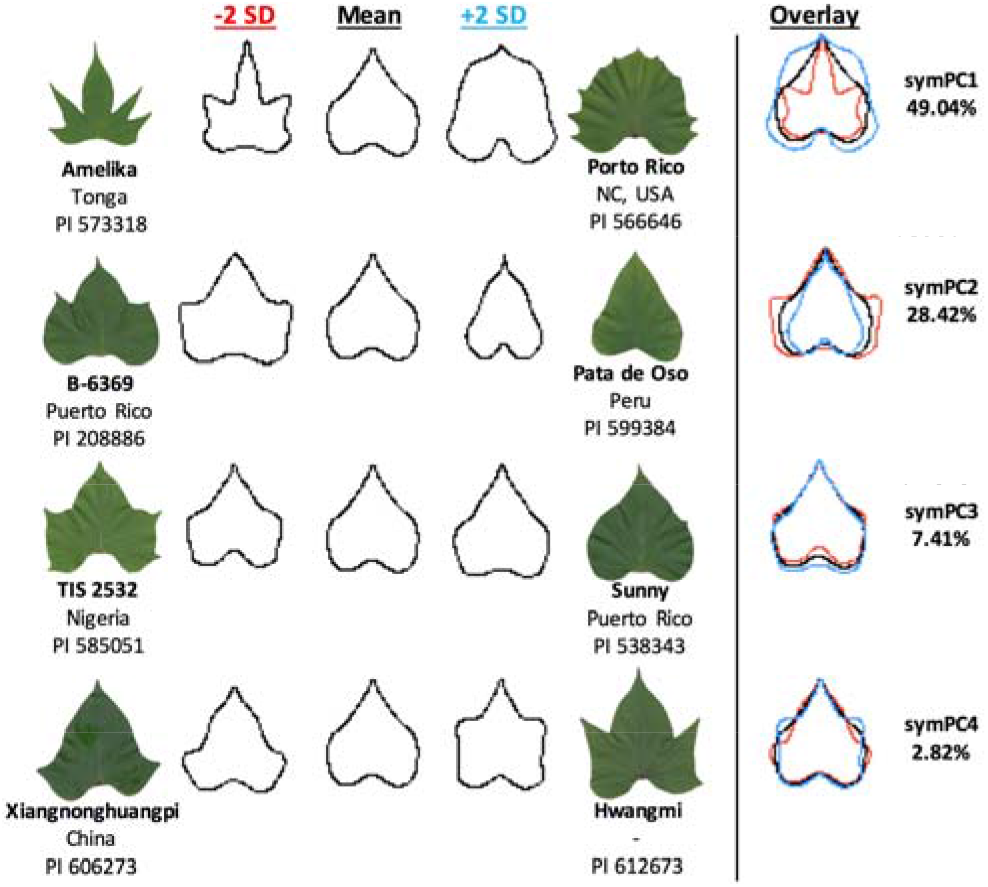
EFDs of symmetrical shape variation. Contours represent eigenleaves resulting from PCA on symmetrical shape (symPC) on EFDs. Shown are the first four PCs with the percent variation explained by each; 87.79% of total variation is explained. −2SD (red) and + 2SD (blue) represent two units of standard deviation from the mean along the PC. Representative leaves of accessions with extreme PC values are shown.

Further, we calculated correlation matrices for traditional shape descriptors and EFD symPCs to determine if they capture different aspects of leaf shape (Fig. S2). We found that symPC1 is correlated with circularity (r = 0.20; P = 0.03) and solidity (r = 0.20; P = 0.02), which is expected as symPC1 partially captures shape differences due to lobing. Additionally, circularity was highly correlated with solidity (r = 0.96; P < 0.001). This is not surprising as circularity is a measure of serrations and lobing whereas solidity is a measure of deep lobing; leaves having deep lobes (and lacking serrations) will thus have similar values of circularity and solidity.

### Sequencing and *de novo* assembly of *I. batatas* transcriptome

We performed a transcriptomic survey to identify gene expression changes associated with the leaf shape traits described above. For our analyses of the transcriptome, Illumina HiSeq2500 returned a total of 266 million (125bp) paired-end sequence reads; on average, each individual had 14 million (M) reads (GEO Submission ID-GSE128065) which was used to construct a *de-novo* transcriptome assembly (sequence statistics are presented in Table 2). The results from BUSCO (Simão *et al*., 2015) indicate that the *de novo* transcriptome assembly is of high quality with 91.32% (1315/1440) complete genes found (single copy genes ~87%) of which only 4.51% were duplicates. Additionally, only 6.32% of genes were missing from the assembled transcriptome. Thus, our sequencing and assembly strategy produced a relatively complete transcriptome. Using blastx, 24,565 transcripts were annotated by the functional description of their top 20 hits. The transcriptome is available at Transcriptome Shotgun Assembly Database hosted by NCBI (TSA accession # GHHM01000000).

**Table 2.** Sequence statistics of the reference transcriptome obtained from EvidentialGene pipeline.

| Number of transcripts | Min Len (nt) | Max Len (nt) | Number of bases | Mean Len (nt) | ORF percent | n50 (nt) | % reads mapped |
| --- | --- | --- | --- | --- | --- | --- | --- |
| 33,684 | 200 | 16,428 | 35,769,411 | 1,062 | 79.95% | 1,608 | 77% |

### Identification and functional annotation of differentially expressed transcripts (DETs)

As a first step towards understanding the genetic control of leaf shape, we identified gene expression changes associated with multiple leaf shape traits -- circularity, aspect ratio (latitudinal expansion) and the symPCs obtained from the EFD analysis. We did not consider solidity and symPC4 due to their high correlation to circularity and low level of variation captured, respectively. On average, we found that 11 million unique paired-end reads per individual (range 7.66M – 14.23M) mapped back to the reference transcriptome (net mapping efficiency of 89.65% with the paired-end high-quality reads). This indicates that we had sufficient read depth (>10M) to continue with our differential expression analysis (as shown by Wang *et al*., 2011).

We uncovered 530 DETs associated with our leaf shape traits (Figure 5, Table S2). Specifically, we found 47 DETs associated with circularity, and 158 DETs associated with aspect ratio. For the symPCs examined, we found 121 DETs associated with symPC1, 148 DETs with symPC2 and 56 DETs with symPC3. Functional annotation of these DETs uncovered putative leaf shape genes (Table 3). As an example, for circularity, FAR1-related sequence 5 (or *FRS5)*, a putative transcription factor involved in regulating light control of development, is differentially regulated with log fold-change of 5.77. Among other DETs for circularity, we found genes that are involved in regulating cell proliferation and organ morphogenesis (EXO70A1-like and extra-large guanine nucleotide-binding protein) and could be involved in regulating leaf dissection.

**Figure 5:**
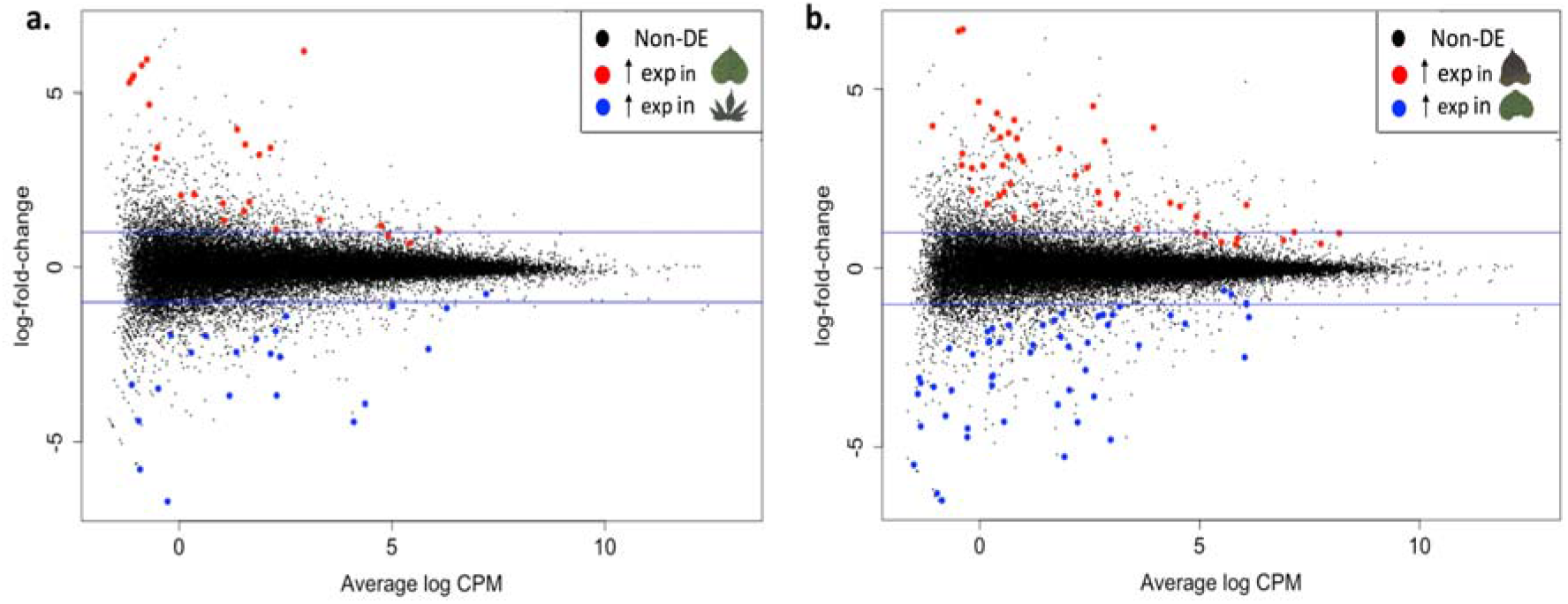
Plot of log-fold change against log CPM (counts per million) with differentially expressed transcripts highlighted (red and blue dots), **a.** Red and blue dots represent transcripts with higher expression in entire and lobed respectively, **b.** Red and blue dots represent higher expression in high aspect ratio and low aspect ratio individuals respectively.

**Table 3.** Candidate genes maintaining variation in leaf traits (circularity, AR and symPCs) identified from the set of differentially expressed transcripts (DETs) in *Ipomoea batatas*.

| Transcript ID | LogFC | FDR | Gene Description |
| --- | --- | --- | --- |
| **Circularity** |  |  |  |
| trn22514 | 5.77 | 0.003 | FAR1-RELATED SEQUENCE |
| trn27202 | 2.08 | 0.021 | Exocyst complex component EXO70A1 |
| trn24081 | 1.33 | 0.033 | Extra-large guanine nucleotide-binding protein |
| **Aspect Ratio** |  |  |  |
| trn9778 | -2.95 | 0.035 | Chalcone Synthase (CHS) |
| trn24267 | 2.55 | 0.00 | Feruloyl CoA 6'-hydroxylase 2 |
| trn25053 | -1.85 | 0.021 | Protein LIGHT-DEPENDENT SHORT HYPOCOTYLS 10 |
| **symPC1** |  |  |  |
| trn27227 | 1.56 | 0.018 | Homeobox-leucine zipper HAT22 |
| trn23566 | 3.54 | 0.00 | FAR1-RELATED SEQUENCE 7 |
| **symPC2** |  |  |  |
| trn27049 | -3.09 | 0.009 | Chalcone Synthase |
| trn28352 | -3.52 | 0.00 | Chalcone Synthase |
| trn9093 | -2.21 | 0.00 | Sporamin B |

Among the 158 transcripts differentially expressed for AR (broad leaves vs rounder leaves), two genes have been shown in literature to alter the longitudinal vs latitudinal expansion of the leaves. These are *CHS* (chalcone synthase), an enzyme involved in the production of chalcones involved in flavonoid biosynthesis, and feruloyl CoA 6’-hydroxylase which is involved in scopoletin biosynthesis and causes post-harvest physiological deterioration in cassava (Liu *et al*., 2017). Finally, we also found LIGHT-DEPENDENT SHORT HYPOCOTYL 10 *(LSH10)*, to be significantly downregulated (log-fold change of −1.85; P-value < 0.001)

Individuals with extreme values of symPC1, a trait differentiating leaf shape based on lobing and prominence of tips and petiolar sinus, were also analyzed for DETs. Of the 121 transcripts showing differential expression, two genes had interesting functional annotations. We found a homeobox gene *(HAT22)* to be upregulated in individuals with high symPC1 (leaves lacking lobes with flattened leaf tips and enclosed petiolar sinus), with a log-fold change of 1.56. We also found another member of the *FRF1* family -- FAR1-related sequence 7 (or *FRS7)* -- to be upregulated in the high symPC1 individuals, like in the case of circularity.

We found a total of 148 DETs for symPC2, which explains variation in leaf shape due to the differences in the broadness and lobing of the leaf. Again, we found two copies of chalcone synthase *(CHS)* were negatively regulated in high symPC2 individuals. We also found Sporamin B transcript, a tuberous root protein (Yeh *et al*., 1997), to be significantly downregulated (with log-fold change of −2.76; P-value < 0.001). Finally, we identified 56 transcripts that were differentially expressed with respect to symPC3; however, functional annotation revealed that most genes belonged to chloroplastic or mitochondrial genes.

### Field experiment

We performed a field experiment to examine leaf shape in different environments, with the specific goal to determine the extent to which genotype, environment, and GxE altered leaf shape. We found significant variation among accessions (indicating genotypic or genetic variation) for circularity, aspect ratio and solidity (F_73_ = 18.06, F_73_ = 4.22, F_73_ = 21.09; P < 0.001), with accession explaining 73.23%, 38.40% and 77.18% of the total variation, respectively (Table 4). This high variance explained for circularity and solidity is reflected in high heritability values (Table 5; H^2^_MI_cir_= 0.79, H^2^_OH_cir_= 0.73; H^2^_MI_solidity_= 0.82, H^2^_OH_solidity_= 0. 76). We also found evidence of significant block effect (F_8_ = 3.01, P = 0.002; η^2^ = 1.33%) for circularity, whereas aspect ratio and solidity were not significantly influenced by block effects. Garden differences between OH and MI contributed 1.93% (F_1_=15.55, P <0.001) of the variability in AR while the accession by garden interaction contributed 12.95% (a significant GxE effect: F_69_ = 5.01, P = 0.009). AR also had lower heritability within each garden (Table 5; H^2^_MI AR_= 0.39, H^3^_OH AR_= 0.26). Circularity and solidity were not significantly altered by environment and had no significant differences due to GxE.

**Table 4.**
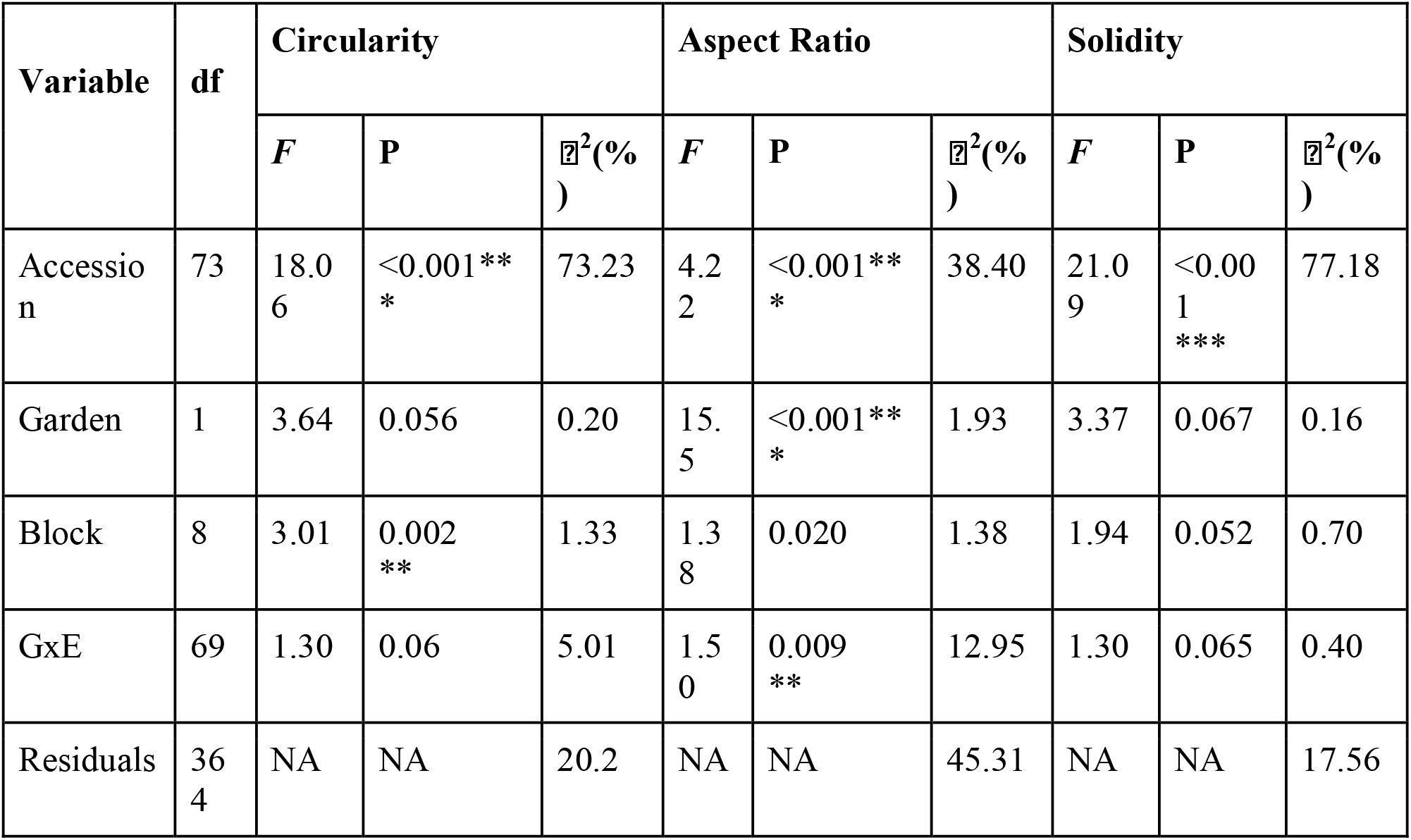
ANOVA table of the leaf shape traits model showing significant explanatory variables. df: degrees of freedom; *F:* value of F-statistic; P: p-value; *η^2^:* eta-squared value.

**Table 5.** Broad-sense heritability values for leaf shape traits in differing environments.

| Env | $H^2$ | | | | | | |
| --- | --- | --- | --- | --- | --- | --- | --- |
|  | Circularity | Aspect Ratio | Solidity | symPC1 | symPC2 | symPC3 | symPC4 |
| MI | 0.79 | 0.39 | 0.82 | 0.80 | 0.58 | 0.70 | 0.69 |
| OH | 0.73 | 0.26 | 0.76 | 0.59 | 0.67 | 0.63 | 0.47 |

\* Note: We can not compare heritability values for EFD symPCs between MI and OH because the expression of traits vary between environments, and hence what the symPCs capture differs between the two environments.

We also examined symmetrical leaf shape variation in both field sites by performing an EFD analysis (Figure 6). EFDs from MI captured variation in leaf shape homologous to the symPCs estimated from greenhouse grown individuals. There was general congruence in symPCs between greenhouse and field grown leaves in MI (i.e., MIsymPC1 (field) ≈symPC1 (greenhouse)), but leaf shape variation captured by EFDs from OH differed significantly in their order of variation explained (Fig. S3). OHsymPC1 explained leaf shape variation due to differences in the broadness and lobing of the leaf (similar to MIsymPC2), whereas OHsymPC2 explained variation due to lobing, tip and petiolar sinus differences (similar to MIsymPC1). This indicates that in OH the majority of leaf shape diversity is primarily due to the broadness of the leaf and secondly due to leaf lobing, while in MI, it is the opposite-- the majority of leaf shape diversity is due to the leaf dissection rather than leaf width. Thus, although traditional shape descriptors are only slightly influenced by the environment, leaf shape as a whole can be altered significantly by the environment.

**Fig. 6.**
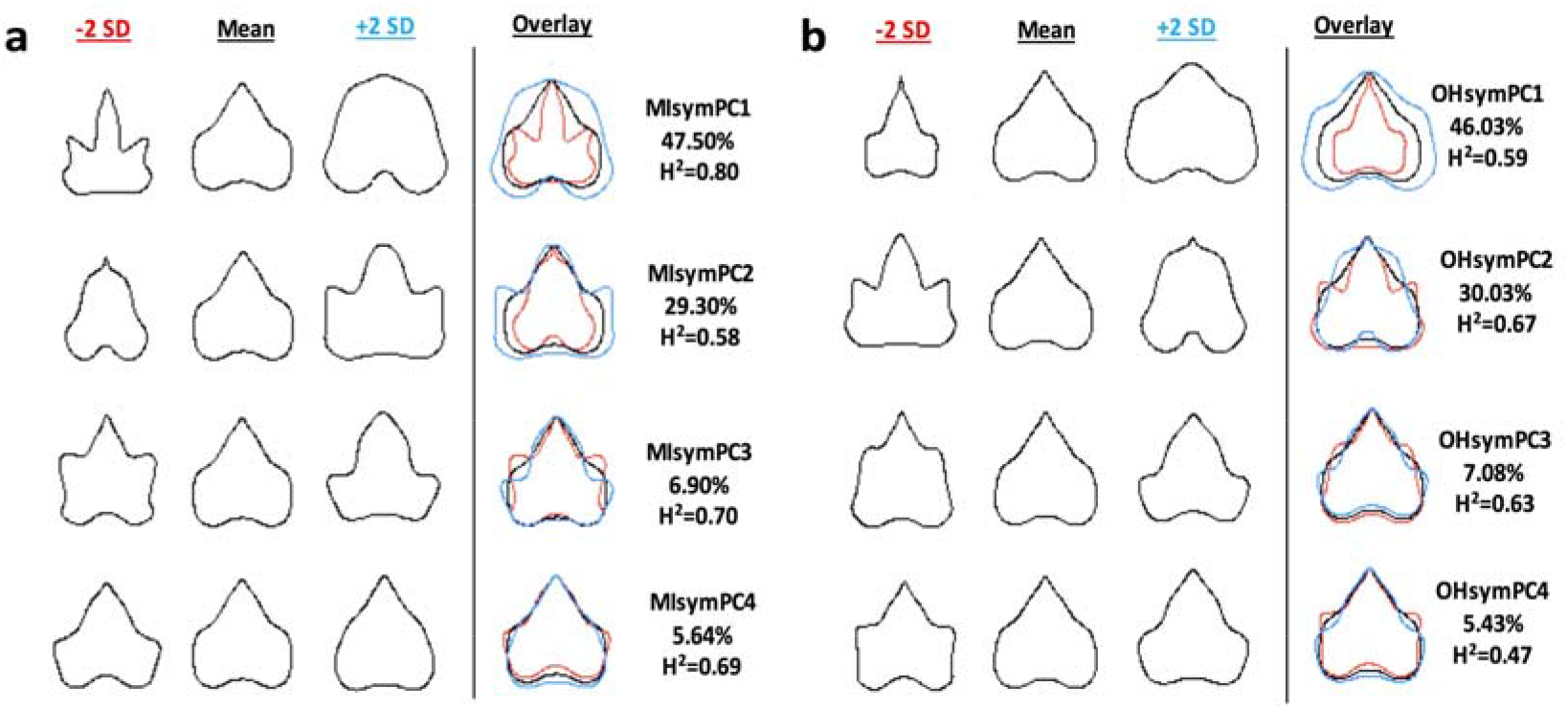
EFDs of symmetrical leaf shape variation among 68 accessions of sweetpotato in the common gardens in Michigan (a) and Ohio (b), respectively.

We also calculated broad sense heritability values for the symPCs in their respective environments and found that H^2^ values ranged from 0.47-0.80 across the symPCs (Figure 6). Heritability values in the OH garden were consistently lower than in the MI garden due to reduced genetic variance and increased environmental variance. Overall, the high heritability values indicate that leaf morphology is controlled to a great extent by genetic factors.

## DISCUSSION

In this study, we examined the extent of leaf shape variation within an agronomically important species, determined the role of genetics, the environment and GxE in altering leaf shape traits, and identified potential candidate genes associated with multiple leaf shape traits. We found evidence of extensive intraspecific morphological variation, with shape differences due to lobing, length-to-width ratio of leaves and the prominence of tip and petiolar sinuses explaining the majority of the variation. We also found that leaf shape has a strong genetic basis with most phenotypic variation attributed to accessional variation, with low or limited influence of GxE. Strikingly, we show that although traditional shape descriptors are only slightly influenced by the environment in this species, when measured comprehensively, leaf shape can be significantly altered by the environment (evident by the change in symPC1 across the MI and OH gardens). Below, we expand on each of our findings, and place them in the context of current knowledge about leaf shape diversity at a species-level as well as what is known about the environmental influence on leaf shape in other species.

### *High morphological diversity of leaf shape in* I. batatas

A recurring question among plant morphologists is the extent to which leaf shape varies among genotypes in a species. This study quantified leaf shape variation among multiple replicated accessions of sweetpotato and identified traits contributing most to leaf shape variation. We focused our morphometric study on three traditional shape descriptors (circularity, aspect ratio and solidity) and then expanded into the more comprehensive Elliptical Fourier Descriptor (EFD) measures.

In our analysis of traditional measures, circularity was found to be the most variable whereas aspect ratio was found to be least variable. Further, the first two principal components of the EFD analysis together accounted for 77.46% of the total variation in leaf shape, and described variation associated with petiolar sinus, tips, and positioning of lobes. Additionally, lack of correlation between symPCs and traditional leaf shape metrics suggests that they capture different features of shape. Only symPC1 was slightly correlated with circularity and solidity. This is not surprising since symPC1 captures variation in leaf shape due to lobing, tip and sinus. No other traits were found to be correlated. Thus, variation captured by the EFD symPCs would have been missed by simply quantifying traditional shape descriptors, suggesting that the use of comprehensive morphometric techniques can help quantify the full extent of shape variation across species. Further, combining the results from traditional morphometric approaches with EFDs revealed that variation in leaf dissection (circularity and symPC1) contributes most to the morphological variation in leaf shape in sweetpotato (Fig. 3 and Fig. 4), similar to that seen in grape (Chitwood *et al*., 2014b). In addition, aspect ratio explains a significant proportion of the remaining variation, unlike in tomato and apple where aspect ratio is the primary trait of variation in leaf shape (Chitwood *et al*., 2013; Migicovsky *et al*., 2017). This indicates that leaf shape variation does not follow a trend across species which is likely due to multiple independent evolution of leaf shape across phylogenetic taxas (Nicotra *et al*., 2011).

### Gene transcripts underlying leaf shape variation

To further our understanding of gene expression changes underlying leaf shape diversity, we sequenced transcriptomes of 19 accessions and assembled a high-quality gene expression database for performing a differential expression analysis in *I. batatas*. We found 47 genes that were differentially expressed for circularity and 121 DETs for symPC1 -- a trait that accounts for leaf shape differences due to leaf dissection, prominence of the tip and petiolar sinus. Functional annotations of these genes identified potential candidates that could contribute to leaf shape dissection in *I. batatas* (Table 3). The most promising candidate is *FRS*gene; we found *FRS5* and *FRS7* to be upregulated in non-dissected individuals in the differential analysis for circularity and symPC1, respectively. *FRS* is a putative transcription factor and contains the DNA binding domain needed to bind the RB-box promoter region of *STM (SHOOT MERISTEMLESS)* (Aguilar-Martínez *et al*., 2015), a protein required for leaf serrations (Kawamura *et al*., 2010). *FRS* might bind to *STM* thus regulating its expression. However, we did not find *STM* to be differentially expressed in our datasets. This might be due to no real expression differences or it might indicate that the expression differences is really small and thus the gene is not detected to be differentially expressed.

Furthermore, genes containing homeobox domains have been shown to be associated with leaf dissection in multiple species *--e.g., PTS* in tomato (Kimura et al., 2008), *STM* in *Arabidopsis (Piazza et al., 2010), RCO* in *C. hirsuta* and other Brassicaceae (Vlad *et al*., 2014; Sicard *et al*., 2014) and *LMI1* in cotton (Andres *et al*., 2016). Most of these genes are differentially regulated in the SAM (shoot apical meristem) and P0 (the youngest primordium) to determine the extent of leaf dissection and complexity for the genotype. However, we did not find any homeobox domain containing genes to be differentially expressed in sweetpotato accessions that varied for circularity *(i.e*. lobed vs entire) (Table S3) but found a homeobox leucine-zipper protein *(HAT22)* to be upregulated for high symPC1 individuals. This mismatch could represent a caveat to our transcriptomic sampling stage (P4-P6), which is past the leaf dissection morphogenic stage of development. Thus, although preliminary, our data indicate that the degree of lobing in *I. batatas* might be maintained in later stages of leaf development (P4-P6) by the action of a gene containing a homeobox domain and that the difference in expression required might be very small.

Further, we found a total of 158 differentially expressed genes associated with aspect ratio and 148 DETs associated with symPC2 (leaf shape due to the differences in the broadness and lobing). Based on the function of the homologs of these genes, we identified promising putative candidate genes responsible for broad leaved phenotypes (Table 3). In apples, a transgenic *CHS* silenced individual developed longer leaves when supplied with naringenin, thus altering leaf AR. This indicates that higher expression of *CHS* (and thus naringenin) is responsible for the longitudinal expansion of the leaves and thus downregulation of *CHS* could lead to broader leaves due to the lack of longitudinal expansion. Another gene of interest that we found differentially expressed for aspect ratio, feruloyl CoA 6’-hydroxylase, produces broader leaved phenotypes of cassava when silenced (Liu *et al*., 2017). Interestingly, however, we found *higher* expression of feruloyl CoA 6’-hydroxylase2 in broader-leaved, compared to the rounder-leaved individuals. Finally, the differentially expressed *LSH10* belongs to the family of *LSH* genes, which have been shown to interact with BOP (BLADE-ON-PETIOLE) and regulate *PTS* (PETROSELINUM) expression, a gene that regulates *KNOX* genes, and thus leaf complexity (Ichihashi et al. 2014). This indicates the potential role of *LSH* gene in regulating both leaf broadness and complexity in this species.

### Factors influencing leaf shape traits in multiple environments

While studies often examine the potential for plasticity in leaf shape traits (McLellan, 2000; Royer *et al*., 2009; Viscosi, 2015), the relative influence of genetic background, environment and gene by environment interactions are less commonly examined. We show that leaf shape traits (circularity, aspect ratio and solidity) in sweetpotato are influenced by multiple effects. Variation in circularity and solidity were mostly attributed to accession (or genotype) and showed little to no effect due to environment or gene by environment interaction. Circularity and solidity have exceptionally high broad-sense heritability values in *I. batatas* (0.76 and 0.79 respectively, averaged between gardens). These traits have likewise been shown to be highly heritable in tomato with heritability values being 0.65 and 0.67, respectively (Chitwood *et al*., 2013). The high PCV for circularity and solidity in *I. batatas* (22.61% and 11.85%) along with high broad-sense heritability indicates that there is a lot of standing variation for these traits that can be actively selected for (or against) by breeders. Furthermore, the lack of plasticity and GxE demonstrate the stability of these simple leaf shape descriptor traits, at least in the environments tested.

Contrary to our results, multiple studies have found that leaf dissection--captured here by our measure of circularity--is a plastic trait that responds to changes in temperature. For example, Royer and colleagues (2009, 2012) found that leaves of *Acer rubrum* were more dissected when grown in cooler environments as compared to warmer environments. A similar trend was observed in grapevine (*Vitis* spp.) (Chitwood *et al*., 2016). However, we found that leaf dissection in sweetpotato is not influenced by the environment. This could reflect that our gardens were not different enough to lead to plastic responses in these two measures of leaf shape. The Ohio garden was consistently warmer (by 2°C on average) and experienced less precipitation than the Michigan garden--the difference between the two gardens was 662.43 mm/month on average throughout the growing season. Although there were environmental differences between gardens, before we conclude that circularity in *I. batatas* is not strongly environmentally responsive, multiple studies in environments that range more widely for temperature will need to be performed.

Comparatively, we found significant variation in aspect ratio due to environment and GxE, explaining 1.93% and 12.95% of the total observed variation in this measure of leaf shape, respectively. This is reflected in the significant alteration of trait values between environments. There was small yet significant differences observed (P < 0.001; 95% CI = 0.009-0.03) between gardens, with clones grown in Michigan consistently showing less round, more elliptical leaves than clones grown in the Ohio garden. However, we still found that 38.40% of the variation in the trait was due to accessional variation which was also indicated in the estimated heritability value of the trait (h^2^ = 0.24). Aspect ratio has been found to be a major source of leaf shape variation in apples and tomatoes with high heritabilities of 0.75 and 0.63, respectively (Chitwood *et al*., 2013; Migicovsky *et al*., 2017). In contrast, we found that this important leaf shape trait is globally not as variable in sweetpotato (4.76% PCV), but it still presents a selection potential. The considerable effect of GxE on aspect ratio indicates that this trait has a genetic component that interacts with the environment leading to varied values between environment.

Further, comparing leaf outlines between two environments, we found that although the traits explaining leaf shape variation are homologous between the two environments, these traits vary in the percent of variation they explain. The heritability of EFD symPCs measured in MI and OH were found to be very high, yet the changes in the amount of variation they explain in their respective environments indicates a strong environmental (and/or GxE) influence on EFD symPCs measured. Although traditional shape descriptors were only slightly controlled by the environment (aspect ratio), we found that the more comprehensive measure of leaf shape can be altered significantly by the environment. This further signifies the importance of measuring leaf shape using methods apart from traditional shape descriptors in multi-environment conditions.

Overall, this work highlights the extensive natural variation in leaf shape within the globally important domesticate *I. batatas*. More broadly, and considering leaf shape analyses from other, mostly domesticated species, leaf shape variation appears to be species specific -- there is no evidence of a shared trait between species that explains the majority of within-species variation. Additionally, we found that most of the variation in the traditional measures of leaf shape appears to be largely controlled by genetic factors in sweetpotato, with a low proportion of variance in leaf shape attributable to environmental differences between gardens. However, when leaf shape was considered more comprehensively and by the use of leaf outlines, we identified a significant influence of the environment, suggesting that studies relying solely on circularity or aspect ratio to describe leaf shape may not capture the extent to which environmental factors can impact leaf development. This multilevel examination highlights the importance of examining morphological variation at the species-level in multiple environments, and using a range of leaf shape phenotypes to comprehensively understand the mechanistic basis (morphological, molecular and environmental) of leaf shape.

## ACKNOWLEDGEMENTS

We are grateful to Robert Jarrett (USDA, Tifton, GA) for providing the sweetpotato accessions. We thank the staff of Matthaei Botanical Garden and Nichols Arboretum (MBGNA, Ann Arbor) for helping us grow and maintain the accessions. We also thank Dan York, Tyler Marrs, Tilottama Roy, Andrew Fox, Jordan Francisco, Yufei Gao, Abby Singletary and Nicholas Tomeo for growing the plants in the field and collecting data. We are also thankful to Daniel H. Chitwood for comments on the manuscript. Funding for this work was provided by the University of Michigan and Ohio University.

## AUTHOR CONTRIBUTIONS

RSB and DMR conceived of the research idea; SG, RSB and DMR performed the experiments and SG performed data analyses with RSB’s supervision. SG wrote the manuscript in consultation with RSB, DMR, JRS.

## Supporting Information

**Method S1** RNA-Seq data processing and transcriptome analysis.

**Fig. S1** Green-house grown accessions selected for transcriptomic analysis.

**Fig. S2** Correlation plot between leaf shape traits (traditional and EFD PCs).

**Fig. S3** Leaf shape variation captured by EFDs from MI and OH differing significantly in their order of variation explained.

**Table S1** Accession IDs with their source and location of origin used in this study.

**Table S2** Differentially expressed transcripts associated with leaf shape traits found in this study.

**Table S3** Raw read counts of orthologs of homeobox domain genes within the assembled transcriptomes, for accessions chosen for circularity RNA-Seq analysis.

