## Supplementary figures and images for "Assessing the remarkable morphological diversity and transcriptomic basis of leaf shape in *Ipomoea batatas* (sweetpotato)"

### Fig. S1

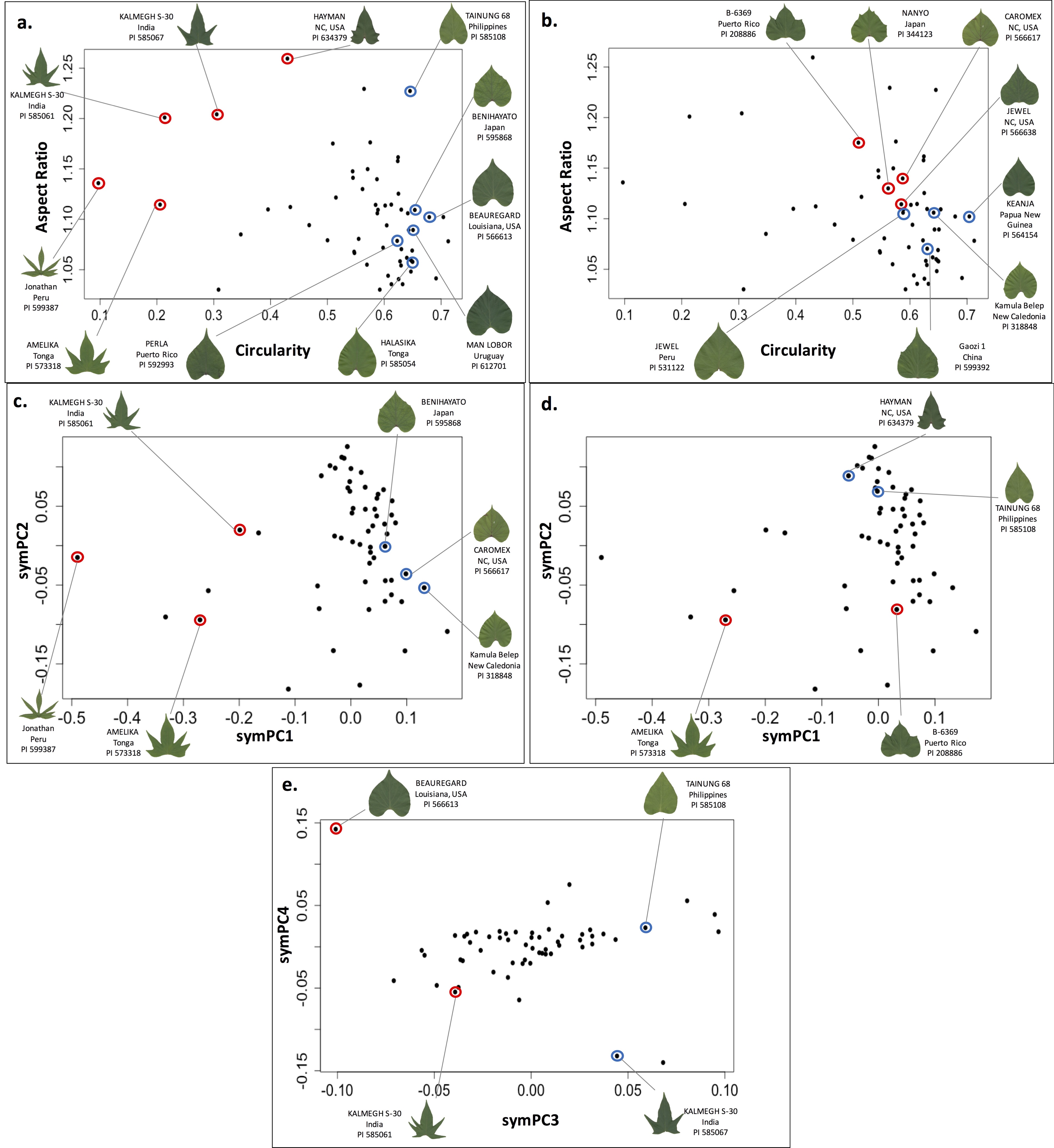

### Fig. S2

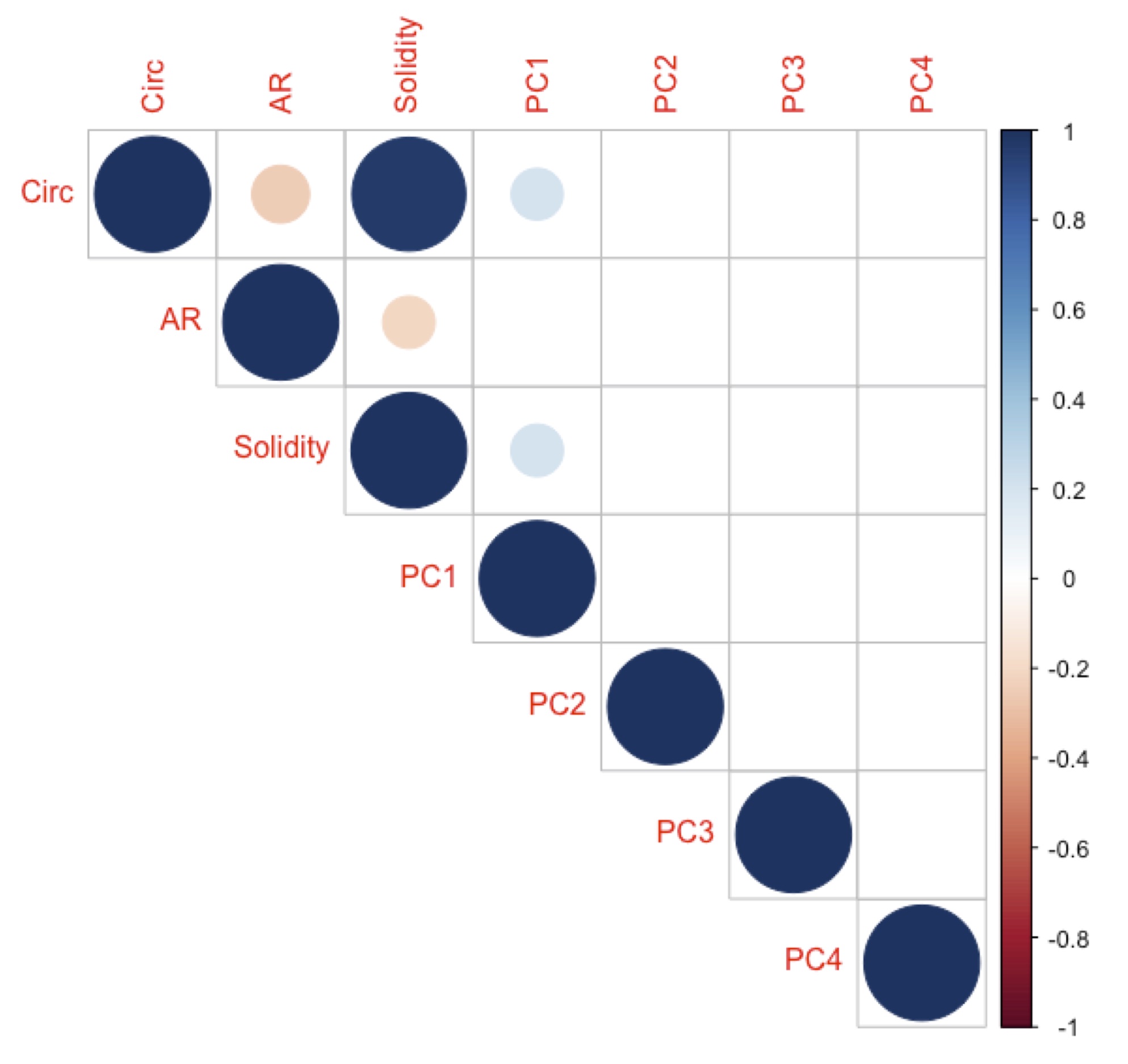

### Fig. S3

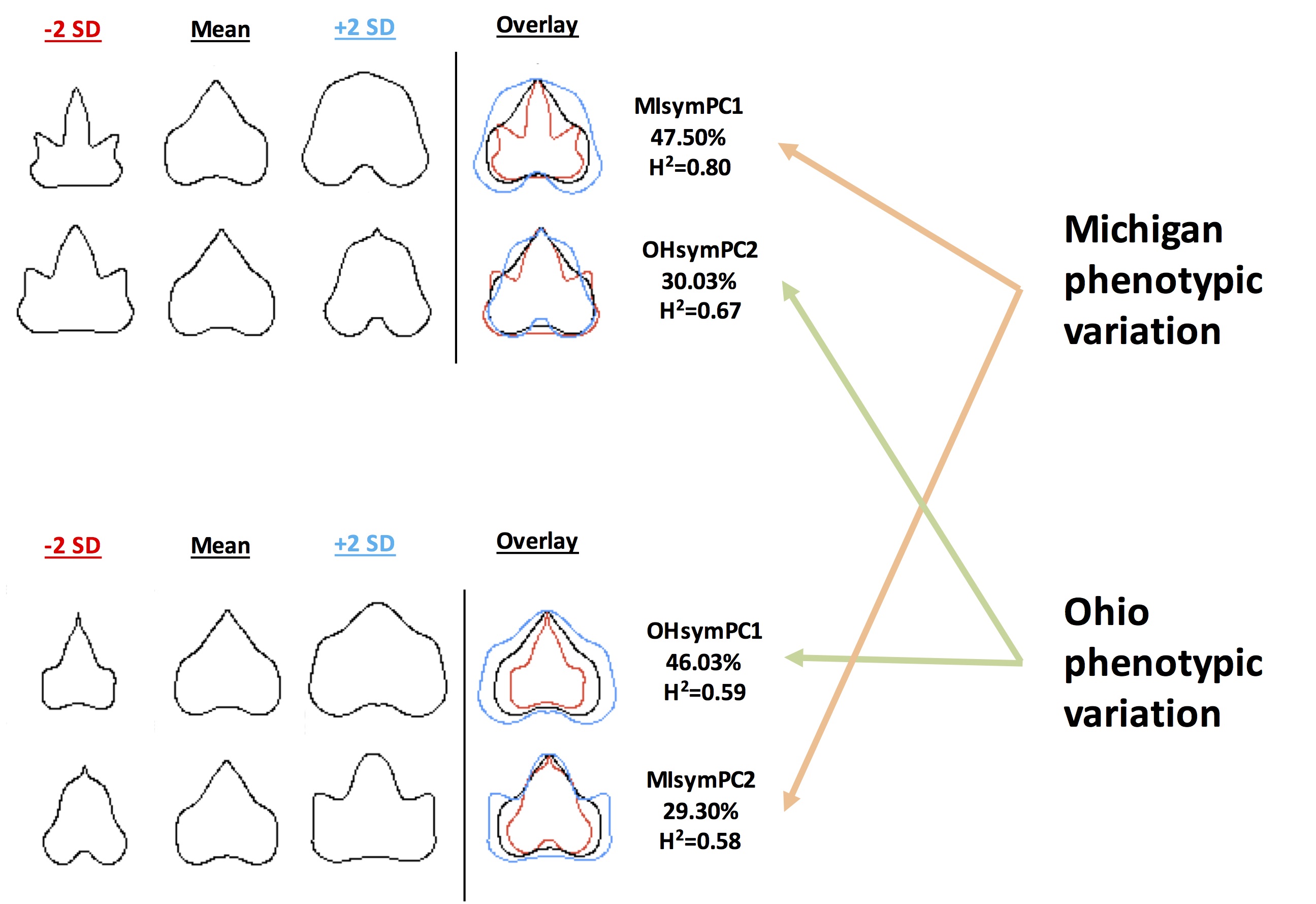
