## Supplementary material for "Assessing the remarkable morphological diversity and transcriptomic basis of leaf shape in *Ipomoea batatas* (sweetpotato)": Method S1

**Method: RNA-Seq data processing and transcriptome analysis**

Briefly, we performed quality control for the obtained raw reads to trim the adaptors, discard low-quality reads and eliminate poor-quality bases. We used cutadapt v1.4 (Martin, 2011) to remove the adapters, and trimmomatic v0.36(Bolger *et al.*, 2014) to clean the reads based on length and quality score. Further, we performed error correction of the RNA-Seq data using rcorrecter(Song & Florea, 2015) to retain only high-quality data.

Next, we used filtered reads separately from one entire and one lobed individual, randomly chosen, for *de novo* transcriptome assembly, which served as a reference transcriptome for differential analysis. To get a comprehensive assembly, we used both a single k-mer approach, using Trinity v2.2.0(Grabherr *et al.*, 2011), with k=25, and multi k-mer approach, using Velvet/Oases v1.2.10(Zerbino & Birney, 2008; Schulz *et al.*, 2012) with k-mer ranging from 23-41 and 93-99 with a step-size of 2. Next, we used the EvidentialGene tr2aacds pipeline (<http://arthropods.eugenes.org/EvidentialGene/trassembly.html>) to merge all the assemblies to remove redundancy and to get biologically most useful set of transcripts.

We then evaluated the obtained set of primary transcripts using TransRate v1.0.3(Smith-Unna *et al.*, 2016) and BUSCO v3- Benchmarking Universal Single-Copy Orthologs(Simão *et al.*, 2015), which reports basic summary statistics (like n50, % reads mapped, etc.) and checks for the completeness of the transcriptome respectively. For annotation of this *de novo* assembled transcriptome, we blasted the transcripts against the NR database with an e-value threshold of 10^-6^ and other default parameters and used only the top 20 hits for annotation. Additionally, we identified conserved protein domains by searching through the InterPro collection of databases. We used the results from these to functionally annotate using BLAST2GO v4.1.9(Conesa *et al.*, 2005) by identification of Gene Ontology (GO) Slim terms and KEGG pathways.
